# The norepinephrine transporter regulates dopamine-dependent synaptic plasticity in the mouse dorsal hippocampus

**DOI:** 10.1101/793265

**Authors:** Alex Sonneborn, Robert W. Greene

## Abstract

The rodent dorsal hippocampus is essential for episodic memory consolidation, a process dependent on dopamine D1-like receptor activation. It was previously thought that the ventral tegmental area provided the only supply of dopamine to dorsal hippocampus, but several recent studies have established the locus coeruleus (LC) as a second major source. However, the mechanism for LC-dependent dopamine release has never been explored. Our data identify norepinephrine transporter reversal as one plausible mechanism by demonstrating that transporter blockade can reduce dopamine-dependent long-term potentiation in hippocampal slices. We also suggest that presynaptic NMDA receptors on LC terminals may initiate this norepinephrine transporter reversal. Furthermore, as dopamine and norepinephrine should be co-released from the LC, we show that they act together to enhance synaptic strength. Since LC activity is highly correlated with attentional processes and memory, these experiments provide insight into how selective attention influences memory formation at the synaptic and circuit levels.

## Introduction

Adrenergic signaling in the mammalian brain is largely controlled by a network of remarkably divergent axon projections arising from locus coeruleus (LC) neurons^1,2^. These LC axons were once thought to exclusively release norepinephrine (NE)^3^, but recent chemical evidence reveals that their specific activation can also increase extracellular dopamine (DA)^4-7^. In accordance with this, LC stimulation is sufficient to modulate DA-dependent changes in learning and synaptic physiology within the rodent dorsal hippocampus^5,8-10^. Dopamine D1-like receptors are abundantly expressed in this region, where they play an essential role in promoting many forms of long-term synaptic potentiation (LTP), especially in area CA1^11^. However, in CA1, projections from canonical DA-releasing nuclei such as the ventral tegmental area (VTA) are sparse compared to those of the LC^8,12^, indicating that DA receptor activation in this area is mainly due to LC activity. Yet despite data supporting the LC as the main source of DA in dorsal hippocampus, the mechanism underlying its release has never been explored.

One hypothesis for the mechanism of LC DA release is by reverse transport through the norepinephrine transporter (NET). Under normal conditions, the NET is responsible for the reuptake of both NE and DA after they are released^13,14^. In contrast, the presence of amphetamines allows the NET to *efflux* catecholamines from LC varicosities^15^, and DA released in this way potentiates synaptic strength in dorsal CA1^16^. Furthermore, the closely related dopamine transporter (DAT) can reverse its flux under more physiological conditions than amphetamine application (for a review, see Leviel (2017))^17^. These conditions include a rise in intracellular [Na^+^] and [Ca^2+^] following action potential firing^18,19^, activation of NMDA receptors^20,21^, and phosphorylation by CAMKII or PKC^22^. Because the amino acid sequences of DAT and NET are almost 80% homologous^23^, we propose that the NET will also efflux cytosolic DA from LC axons under similar physiological conditions. Below we investigate this possibility in the dorsal hippocampus, where DAT expression is not detectable^16,24^, by designing a DA-dependent LTP that is significantly attenuated after the NET is blocked.

In support of a more detailed model for NET-mediated DA release, an existing theory posits that high-frequency glutamate activity may play a role^25^. The authors speculate that elevated pyramidal cell firing in response to environmental stimuli can result in glutamate spillover^26^, leading to activation of presynaptic NMDA receptors on LC terminals and enhanced vesicular NE release. Taking this idea one step further, Olivier *et al.* discovered that an NMDA-dependent rise in striatal DA is nearly abolished by GBR12909, a selective DAT blocker^20^. This indicates that NMDA receptors can somehow interact with transporters and change the direction of catecholamine flux. Comparably, early studies in dorsal hippocampus reported increased extracellular NE and DA after NMDA receptor agonist application^27,28^. To our knowledge, no studies have attempted to associate this DA transmission with NET reversal, or looked at its ability to regulate synaptic plasticity. With this in mind, we deleted NMDA receptors from catecholamine terminals and saw a decrease in DA-dependent LTP in dorsal hippocampus.

Lastly, given that both DA and NE can modulate synaptic plasticity in dorsal CA1^29^, along with the indication of their co-release from the LC, we presume that they are working together to influence synaptic strength in this region^30^. Our final experiment shows that simultaneous application of DA and NE, but not either of them alone, can strengthen a weaker form of hippocampal LTP. The implications of these results are then discussed in the context of the LC’s purported role in selective attention^31^, and how glutamate can interact with catecholamines to organize attention-driven memory formation at the synaptic and circuit-specific levels.

## Results

### The norepinephrine transporter (NET) contributes to dopamine-dependent potentiation in the dorsal hippocampus

If the NET is capable of controlling DA efflux in dorsal hippocampus, then blocking it should attenuate DA-dependent synaptic potentiation. To test this, we developed a strong theta-burst LTP protocol (strLTP) by stimulating CA3 axons and recording the slope of field excitatory postsynaptic potentials (fEPSPs) from stratum radiatum dendrites of CA1 (Fig.1A-D). This protocol was based on previous methods used to generate catecholamine-dependent potentiation in hippocampus^32,33^. Importantly, our strLTP was not blocked by co-application of β-adrenergic (propranolol) and α1-adrenergic (prazosin) receptor antagonists (Fig. 1E, comparison between strLTP with no-drug from Fig. 1D and strLTP with drug from Fig. 2A). However, following the addition of the D1-like receptor antagonist SCH 23390, a robust blockade of LTP occurred over the last 30 minutes of recording (Fig. 2A, red traces), indicating that strLTP is dependent on DA receptors, but not adrenergic receptors. Next, we administered the same strLTP stimulation, but substituted nisoxetine, a NET blocker, for SCH 23390. Treatment with nisoxetine produced a similar reduction in LTP (Fig. 2B, green traces), suggesting that DA signaling in the dorsal hippocampus requires NET activity. Likewise, a genetic deletion of the NET from LC neurons also greatly reduced strLTP amplitude after 1 hour (Supplementary Fig. 1).

**Figure 1.**
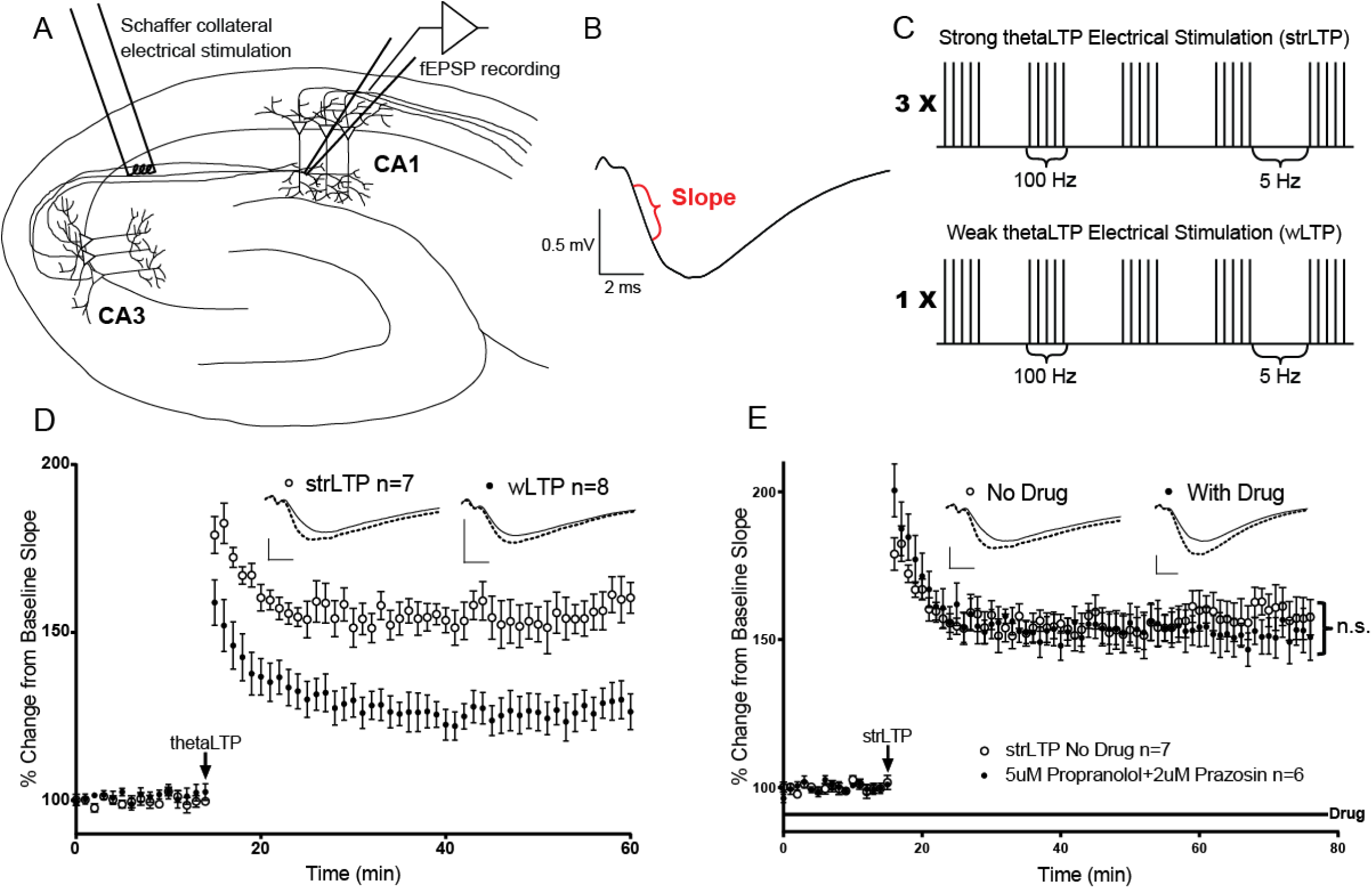
Establishment of weak and strong long-term potentiation (LTP) protocols. **A**, Diagram of a hippocampal slice with electrodes in place. The stimulating electrode (left) is placed in contact with Schaffer collateral axons from the CA3 region about 400 µm from the recording electrode. The recording electrode (right) measures the extracellular field excitatory postsynaptic potential (fEPSP) in stratum radiatum dendrites of CA1. **B**, Example fEPSP from CA1. Data is taken as the initial slope of the voltage trace as shown in red. Scale bars represent the 0.5 millivolt amplitude and two millisecond duration in all of the following figures. **C**, Weak (wLTP, n=8) and strong (strLTP, n=7) Schaffer collateral thetaLTP stimulation protocols (see methods for more details). **D**, Weak versus strong thetaLTP time course. The black arrow represents the moment that either LTP stimulation was given after a 15 minute baseline. Insets are representative traces before and after each stimulation protocol. The solid lines represent an average of baseline traces from 0-15 minutes before LTP stimulation, while dotted lines represent an average of traces from the last 5 minutes of the recording after LTP stimulation. **E**, strLTP (open circles, n=7) is not blocked by the addition of prazosin and propranolol to the bath (closed circles, n=6), F(1, 17) = 0.06472, p=0.8022, ‘n.s.’ stands for ‘not significant’. All data points are represented as mean +/- SEM. Tests for significance were done using a two-way repeated measures ANOVA over the last 30 minutes of strLTP.

**Figure 2.**
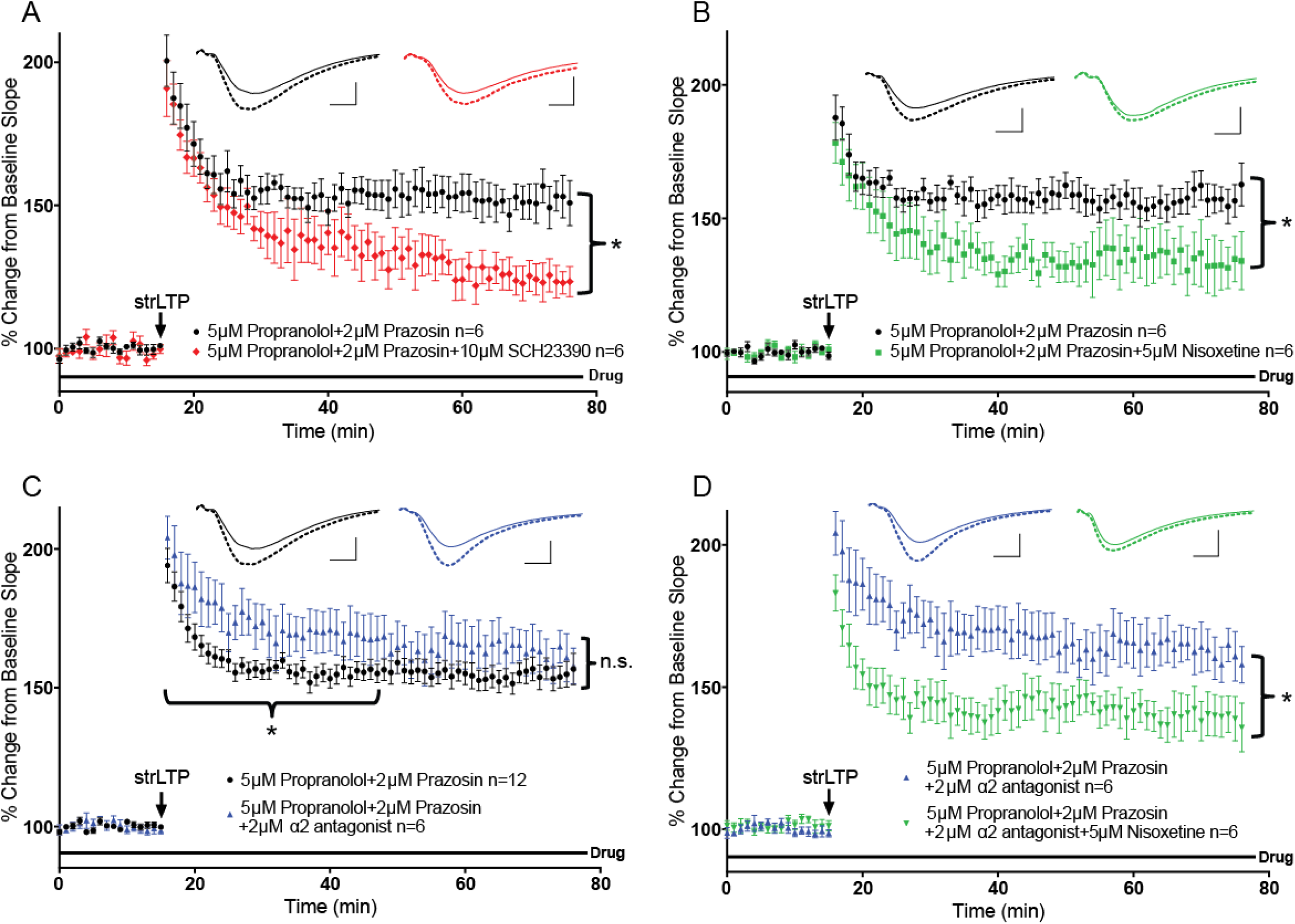
The norepinephrine transporter (NET) contributes to dopamine-dependent long-term potentiation. **A**, The previously established strong LTP protocol (strLTP, black arrow) was not blocked by application of antagonists for β- and α1-adrenergic receptors, propranolol and prazosin, respectively (black circles, n=6, see Fig. 1E). However, application of SCH 23390, a dopamine D1-like receptor antagonist, along with β- and α1 blockers was enough to significantly reduce the last 30 minutes of LTP (red circles, n=6), F(1, 10) = 9.265, p=0.0124. **B**, Similar to **A**, but the D1/5 receptor antagonist was replaced with the NET blocker nisoxetine (green squares, n=6), which was sufficient to attenuate the dopamine-dependent LTP (black circles, n=6), F(1, 10) = 5.028, p=0.0488. **C**, Blockade of ALL adrenergic receptors (by adding an α2 autoreceptor antagonist) selectively increased the *first* 30 minutes of LTP (blue triangles, n=6) compared to β- and α1 blockers alone (black circles, n=12), F(1, 16) = 4.963, p=0.0406. **D**, Even with all adrenergic receptors blocked (blue triangles, n=6) the application of nisoxetine was still able to significantly reduce LTP (green triangles, n=6), F(1, 10) = 5.521, p=0.0407. All data points are represented as mean +/- SEM. Tests for significance were done using a two-way repeated measures ANOVA over the last 30 minutes of strLTP, or during the first 30 minutes as shown in panel C. Asterisks represent p-values <0.05, while ‘n.s.’ stands for ‘not significant’.

### Blocking α2-adrenergic receptors does not reduce the effect of NET antagonism

Because blocking the NET will flood synapses with NE, one possible confound is over-activation of inhibitory α2-adrenergic autoreceptors, leading to a decrease in overall LC excitability and neurotransmitter release^34^. This may cause a reduction in LTP based on an indirect decrease in total NE and/or DA levels. To control for this, we repeated the aforementioned experiments with the inclusion of RS 79948, an α2-receptor antagonist, in the bath with propranolol and prazosin. Interestingly, adding RS 79948 caused a significant increase in fEPSP slope over the first 30 minutes after strLTP stimulation, but not the last 30 minutes (Fig. 2C, blue traces). This effect could be due to greater vesicular NE release reaching concentrations high enough to displace propranolol and activate β-adrenergic receptors, a process known to enhance early LTP^35^. In line with our prior results, the further addition of nisoxetine was still able to diminish the magnitude of strLTP over the last 30 minutes (Fig. 2D, green traces), reinforcing the finding that NET may contribute to DA signaling in dorsal hippocampus.

### NMDA receptor knock-out from catecholamine neurons reduces the magnitude of dopamine-dependent LTP

Activation of glutamate receptors, in particular NMDA, is capable of enhancing catecholamine release in hippocampus^27,28,36^. Expanding on the mechanism of DA release from the NET, we asked if presynaptic NMDA receptors on LC terminals were functionally coupled to NET reversal, and thus involved in our NET/DA-dependent LTP. To expand on the mechanism of DA release from the NET, we asked if presynaptic NMDA receptors on LC terminals could functionally be coupled to NET reversal, and thus involved in our NET/DA-dependent LTP. To approach this question, we first confirmed that NMDA receptors co-localized with the norepinephrine transporter on LC axon terminals in the dorsal hippocampus (Supplementary Fig. 2). To our knowledge, this is the first time that co-localization of these proteins has been shown in LC terminals in any area of the brain.

We next wanted to check if this co-localization was important for the expression or maintenance of LTP in dorsal CA1. To do this, the NR1 subunit of NMDA receptors had to be genetically deleted from catecholamine neurons, since blocking NMDA receptors would prevent LTP. This was done by crossing a mouse expressing Cre recombinase under the control of the tyrosine hydroxylase (TH) promoter with a floxed NMDA-NR1 subunit mouse. Cre-negative controls for these mice showed normal strLTP (Fig. 3, filled circles), whereas the LC-NR1 knockouts exhibited decreased LTP magnitude throughout the full hour after LTP induction (Fig. 3, open circles). Since dorsal CA1 receives very little VTA input, we interpreted these effects as being predominantly due to NMDA deletion from LC neurons. However, the results do not rule out possible compensatory effects of NMDA receptor knockout resulting from Cre expression in TH-positive neurons during development^37^.

**Figure 3.**
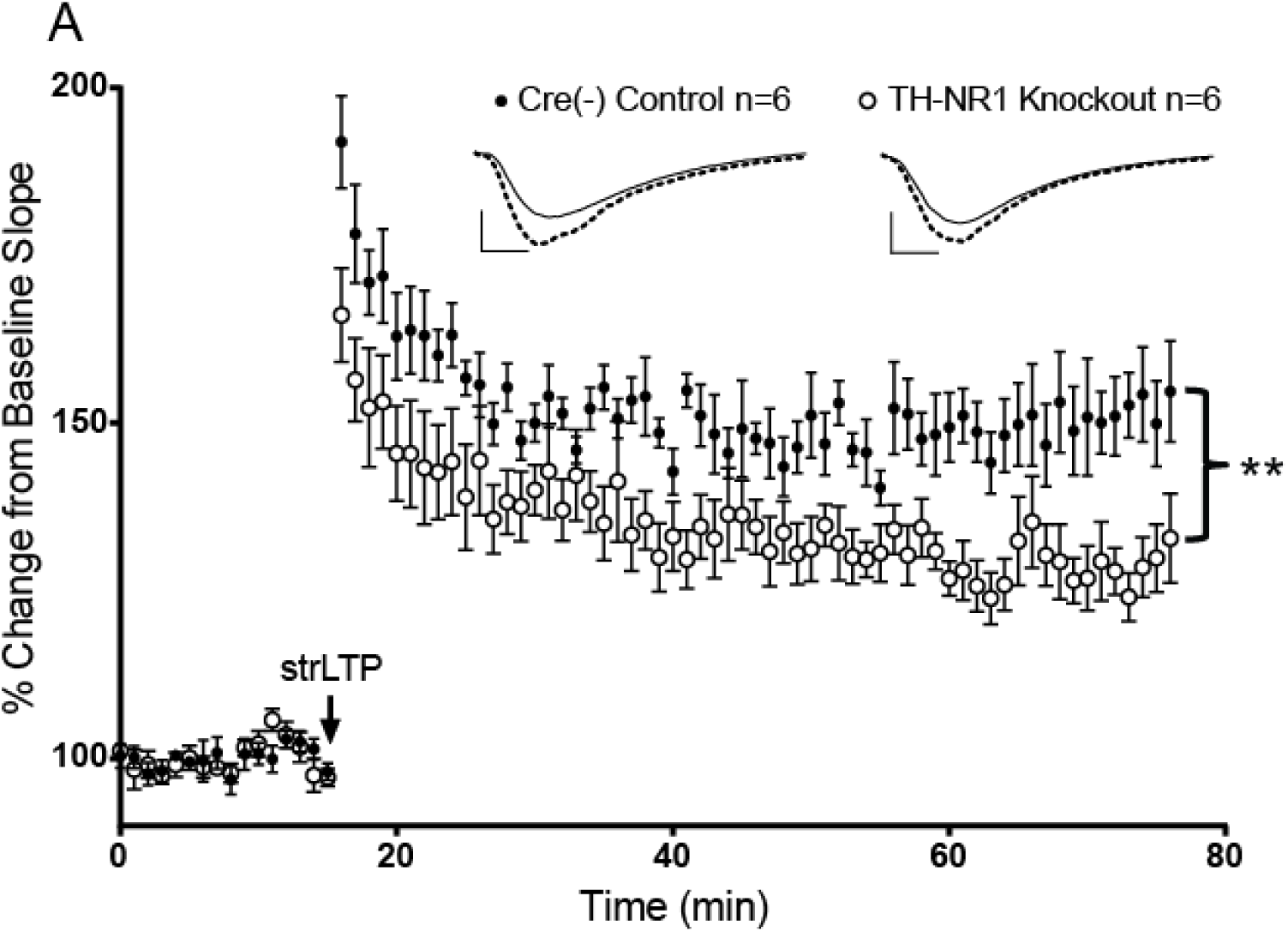
Knocking out NMDA receptors from catecholamine neurons reduces the magnitude of dopamine-dependent LTP in dorsal hippocampus. **A**, The same strong LTP protocol used previously (strLTP, black arrow) was administered in slices from Cre(-) control mice (black circles, n=6) and NMDA-NR1 subunit knockout mice (open circles, n=6). All data points are represented as mean +/- SEM. Tests for significance were done using a two-way repeated measures ANOVA over the last 30 minutes of strLTP. Double asterisk represents a significant difference <0.01, F(1, 10) = 13.24, p=0.0046.

### Concurrent activation of DA and NE receptors is required for LTP enhancement in CA1

The regulation of synaptic plasticity by DA or NE is well documented in dorsal hippocampus^38,39^. For example, D1/D5 receptor antagonists in CA1 can inhibit synaptic potentiation^40^ and contextual learning^41^, while agonists of β-adrenergic receptors are sufficient to lower the threshold for LTP initiation^42^. Yet in dorsal CA1, whether or not coincident activation by both catecholamines is needed to enhance LTP has never been examined. This question remains of great importance, as it has been well established that the LC can release DA and NE together. Accordingly, we developed a weak LTP (wLTP) stimulation paradigm (Fig. 1C&D) and tested if an interaction between DA and NE was necessary to strengthen it.

Bath application of NE alone had no effect on the magnitude of wLTP; although a decrease in baseline glutamatergic signaling before wLTP stimulation was apparent (Fig. 4A, 16-30 mins). The latter phenomenon has been recorded previously^43^, and is likely due to NE activating α1-receptors on interneurons to increase their feed-forward/lateral inhibitory drive^44^. Surprisingly, washing in DA alone had no effect on either wLTP magnitude or basal glutamate transmission (Fig. 4B). This seems counterintuitive considering that activation of D1-like receptors by selective agonists (e.g. SKF-81297) can reliably evoke LTP in dorsal CA1^11^. However, the result is consistent with multiple reports showing no change in CA1 excitatory transmission in response to bath applied DA^45-47^. It is unclear why this occurs, but one explanation could be that over activation of inhibitory D2-like receptors negates the excitatory D1-like receptor activation in this region. Interestingly, even though neither of the catecholamines in isolation was able to produce stronger LTP, their simultaneous application resulted in a significant increase compared to the control wLTP (Fig. 4C). These findings pair well with the following conclusion that the LC utilizes both DA and NE to optimize memory storage, mainly during periods of enhanced attention to salient stimuli.

**Figure 4.**
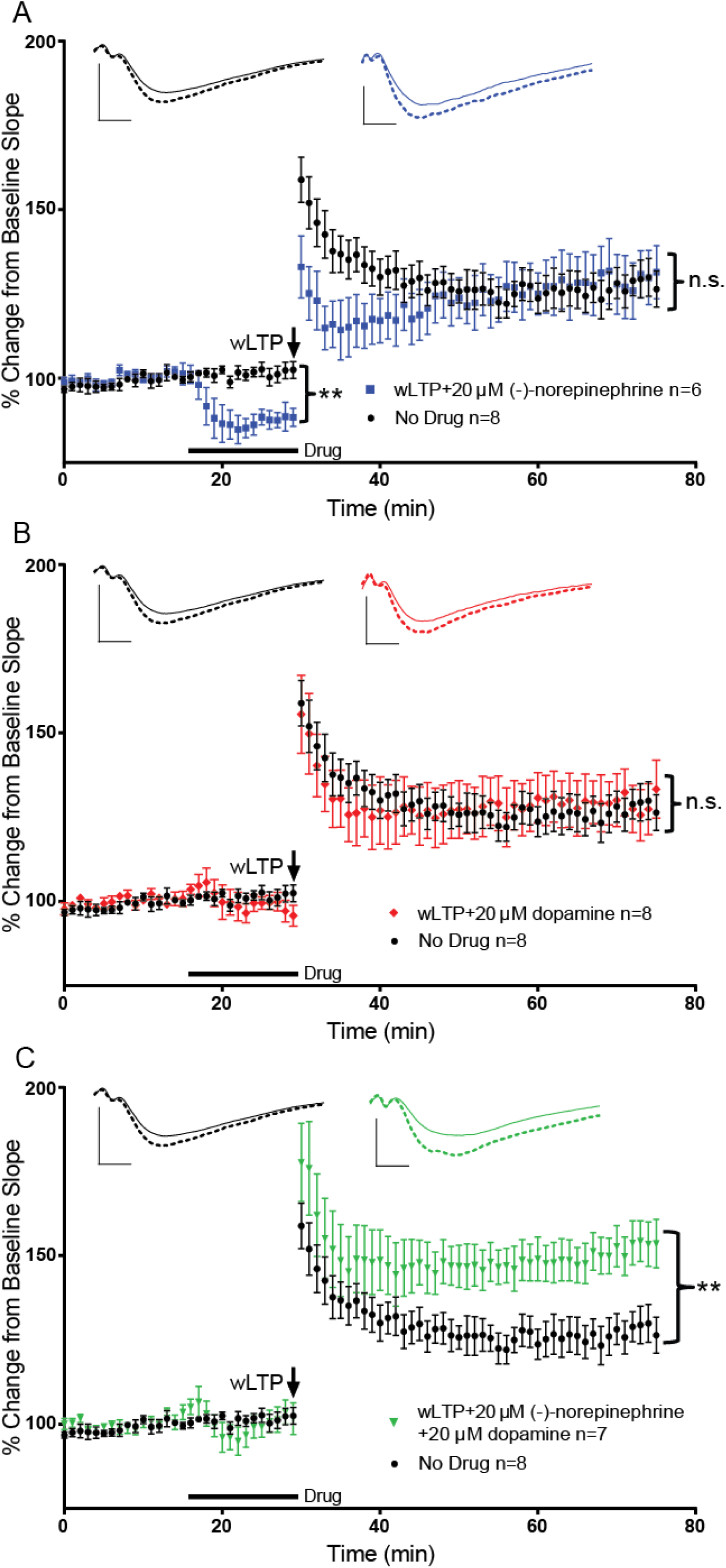
Weak LTP (wLTP) is enhanced by dopamine and norepinephrine together but not by either of them alone. **A**, wLTP (black arrow, black circles, n=8) is not enhanced with bath application of 20 μM norepinephrine alone (blue squares, 61-76 min, n=6), F(1, 12) = 0.05005, p=0.8267. Instead, the application of norepinephrine after a 15 minute baseline significantly reduced the size of the baseline field potential slope (blue squares, 16-30 min, n=6), F(1, 12) = 18.87, p=0.0010. **B**, Bath application of 20 μM dopamine (red diamonds, n=8) was also unable to enhance wLTP, but did not show a similar decrease of baseline slope, F(1, 14) = 0.1202, p=0.7339. **C**, Application of 20 μM norepinephrine and 20 μM dopamine together (green triangles, n=7) produces a significant increase of wLTP, F(1, 13) = 9.318, p=0.0080. All data points are represented as mean +/- SEM. Tests for significance were done using a two-way repeated measures ANOVA over the last 15 minutes of wLTP, or during 15 minutes of drug application as shown in panel A. Asterisks represent p-values < 0.05, double asterisks represent p-values < 0.01, while ‘n.s.’ stands for ‘not significant’.

## Discussion

Taken together, these results allude to the LC orchestrating coincident release of NE and DA in the dorsal hippocampus using two separate mechanisms. The first is the widely accepted vesicular release of NE^48^, and the second is reverse transport of DA from the NET as shown in this study. A reason for separate release mechanisms is still unclear, but one plausible explanation is that they are used to facilitate a molecular link between attention and memory^49^, especially since the LC is heavily involved in both cognitive processes at the behavioral level^50^. Below we postulate that their co-release should only occur when an animal devotes sustained attention to salient stimuli associated with prolonged LC firing. If this happens, NE and DA can interact in CA1 to help exclusively potentiate the most relevant synapses for future memory consolidation (Fig. 4).

When an animal is awake but not experiencing anything particularly interesting in its environment, the LC releases a low, background level of NE from vesicles using a slower, tonic firing pattern (∼3 Hz). In the hippocampus, this leads to a general suppression of activity by activating higher affinity α1-adrenergic receptors on interneurons in the area^43,44^. During times of elevated arousal and selective attention to salient stimuli, higher frequency (∼16 Hz) phasic activity^51^ transiently boosts extracellular levels of NE ^34,52^. These quick increases in NE are theorized to recruit lower affinity β-adrenergic receptors on hippocampal pyramidal cells to amplify more active glutamatergic inputs, while the α1-receptors continue to reduce the noise generated by less active ones^53^. Therefore, NE helps the most immediately relevant and strongest signals prevail over those that are firing slower and likely carrying less important information. Elaborating on this, Mather, et al. ^25^ propose that very active glutamatergic synapses can in turn augment the release of NE after glutamate spillover activates presynaptic NMDA receptors on nearby LC axons. This would create a local positive feedback loop between the most rapidly firing glutamatergic input and phasically active LC terminals, with less active circuits remaining suppressed as they are unable to trigger this positive feedback. In the hippocampus, this system presumably optimizes circuit organization to reduce the overlap between stored memory traces.

Our data expand on these theories and suggest that presynaptic LC-NMDA receptors can similarly initiate DA signaling from LC axons in dorsal hippocampus, since their deletion weakens DA-dependent LTP (Fig. 3). This effect makes sense within the framework of attention being a driving force for memory formation^49^. For instance, when strong glutamatergic signaling in response to salient environmental cues couples with phasic LC firing in CA1, excess glutamate can overflow from the synapse and bind to NMDA receptors on LC terminals^25^. At the same time, salience-guided, phasic action potential firing in LC terminals will influx Na^+^ and Ca^2+^, removing the Mg^2+^ block and allowing even more cation influx through NMDA receptors. Calcium entering the neuron via this process may then promote the function of CAMKII or PKC, kinases capable of interacting with^54^ and phosphorylating transporters to reverse their direction^55^. Likewise, since monoamine transporters are known to move neurotransmitters using the energy stored in Na^+^ gradients^56^, a switch to higher intracellular Na^+^ during action potential bursting could alter the ionic equilibrium enough to move DA out through the NET. In favor of this idea, it is known that the NET can reuptake DA nearly as well as NE in the hippocampus under normal conditions^14^, hinting that the reverse mechanism may be possible. Figure 2 explores this idea and highlights that DA released in this way seems to be physiologically relevant, as blocking or deleting the NET is capable of reducing DA-dependent LTP.

Functionally, this non-canonical DA efflux likely arose as a form of coincidence detection in dorsal CA1. Here it will potentiate only the most prevalent glutamatergic inputs that were selected by the preceding NE modulation of glutamatergic attentional resources. In other words, once a stimulus becomes salient enough to outcompete the background noise, DA is released and interacts with NE to enhance synaptic strength (Fig. 4). This would be necessary to tag specific synapses recruited by the increased glutamate signaling for future memory consolidation, given that DA seems to be more involved in the tagging process than NE^57^. For this reason, having two separate release mechanisms might enable more efficient signal processing and storage of new information, since DA released out of the NET would not interfere with the formation of neural representations driven by vesicular NE release.

Altogether, these observations concerning the NET’s involvement in DA-dependent potentiation are in conflict with a couple of studies that also measured nisoxetine’s effect on LTP in CA1^58,59^. The authors of both papers found no difference in LTP when nisoxetine was present. One explanation could be that, in contrast to our methods, these reports did not use any adrenergic receptor antagonists, potentially leading to excess β-adrenergic receptor recruitment and cAMP dependent LTP enhancement^42^. Although, as mentioned above, our NET knockout assay was also performed in no-drug conditions and produced a massive difference in LTP. A reason for this is not immediately obvious, but we cannot rule out developmental consequences of NET deletion on normal adrenergic system function. However, a critical observation arising from our experiments is the fact that if LC DA was vesicular in origin, then blocking NET should have the opposite effect, as nisoxetine application should lead to an increase in extracellular DA and thus stronger LTP.

In closing, our findings support the idea that the NET and NMDA receptors contribute to DA signaling (Figs. 2&3), and therefore interaction with NE signaling (Fig.4), to regulate attention-guided memory storage in the CA1 region of dorsal hippocampus. One drawback of our methods is that LC fibers were not selectively stimulated. Instead, catecholamine release was elicited by electrical stimulation of all fibers within the range of the stimulating electrode, which could include any other neuromodulatory inputs into CA1 that might interact with the effects of NE and DA (e.g. acetylcholine or serotonin). Also, since we were stimulating with bursts of 100 Hz, this could unnaturally overload LC terminals since their usual maximum firing rate is <20 Hz, leading to DA release out of the NET that would not occur under normal physiological conditions. Future studies may employ specific optogenetic activation of the LC to study this question with greater precision. It will also be necessary to utilize the recently developed genetically encoded fluorescent DA^60,61^ and NE^62^ sensors to probe the dynamics of LC catecholamine co-release in greater detail. In conclusion, although our evidence is indirect, it presents a vital first step towards elucidating the complex interplay between glutamate activity and catecholamine release, not only within the hippocampus, but in all LC terminal fields throughout the central nervous system.

## Methods

### Animal Approval

All animal procedures performed were approved by the animal care and use committee (IACUC) at the University of Texas Southwestern Medical Center and comply with federal regulations set forth by the National Institutes of Health.

### *Ex vivo* slice preparation

Coronal slices (300 µm thick) containing dorsal hippocampus were made from male, wild type, C57BL/6J mice (6-12 weeks old) in low-light conditions to prevent photooxidation of catecholamines. Animals were anesthetized under 1.5-2% isoflurane, after which brains were removed and blocked following rapid decapitation. Slices were prepared using a Leica VT1000S vibratome (Wetzlar, Germany) in ice-cold NMDG ringer solution containing (in mM): 5 NaCl, 90 NMDG (N-Methyl-d-Glucosamine), 37.5 Na-Pyruvate, 12.5 Na-Lactate, 5 Na-Ascorbate, 2.5 KCl, 1.25 NaH_2_PO_4_, 25 NaHCO_3_, 25 Glucose, 10 MgSO_4_.7H_2_0, 0.5 CaCl_2_.2H_2_0. The pH was set between 7.3 and 7.4 using 12 N HCl, the osmolarity was adjusted as needed to ∼315 mOsm using glucose, and the solution was continuously bubbled with 95% O_2_ and 5% CO_2_ gas during slicing. Slices were then transferred and maintained for up to 6 hours, while protected from light, at 30 °C in artificial cerebrospinal fluid containing (aCSF; in mM): 120 NaCl, 3 KCl, 1.25 NaH_2_PO_4_, 1 MgCl_2_, 2 CaCl_2_, 25 NaHCO_3_, and 11 dextrose continuously bubbled with 95% O_2_ and 5% CO_2_ gas.

### Field recordings

After at least 1 hour of recovery in aCSF, slices were transferred to a submersion recording chamber and perfused with aCSF at a rate of 2-3 ml/min at 31-32 °C. Extracellular voltage recordings from the stratum radiatum field of dorsal CA1 were acquired using a borosilicate glass electrode (1-2 MΩ, Sutter Instrument (Novato, CA)) filled with normal aCSF. A bipolar stimulating electrode (FHC, Inc. (Bowdoin, ME)) was also placed in the stratum radiatum of CA1 within ∼400 µm of the recording electrode (see Figure 1A), and stimulus strength was controlled with a stimulus isolator unit (World Precision Instruments, Sarasota, FL). Stimulus strength was set to produce a baseline excitatory field postsynaptic potential (fEPSP) slope (Figure 1B) that was ∼50% of the slope measured following the first appearance of a population spike. This method led to a typical baseline stimulation current of 20-30 μA, while stimulus duration was set to 0.2 ms. Schaffer collateral stimulation was given once every 30 seconds and the average of every two consecutive stimuli was taken. For the DA and NE synergy experiments, a stable 15 minute control baseline was obtained, followed by another 15 minute baseline with drug washed in. At the end of the 15 minute drug wash, a weak theta-burst tetanus was applied consisting of 5 bursts (given at 5 Hz), with each burst containing 5 spikes at 100 Hz (25 total spikes). Baseline stimulation then resumed as described above for 45 minutes. For the NET blockade experiments, the entire experiment was run in the presence of various antagonists (indicated in figures). A 15 minute baseline was obtained, followed by a strong theta-burst tetanus containing 15 bursts (given at 5 Hz), with each burst containing 5 spikes at 100 Hz (75 total spikes). Baseline stimulation then resumed as described above for 60 minutes. For Supplementary Data 1, a different LTP stimulation protocol was used that consisted of a 1 second long train of 100 Hz (100 total spikes), without any antagonist application. All experiments were performed in low-light conditions to avoid photooxidation of catecholamines. Data was acquired using a Multiclamp 700B amplifier and pCLAMP 10 software (Molecular Devices, San Jose, CA). The signal was low-pass filtered online at 2 kHz using the Multiclamp 700B Commander software, and then digitized at 20 kHz using a Digidata 1440A (Molecular Devices, San Jose, CA).

### Staining and imaging

Dorsal hippocampal sections, 30 µM thick, were cut with a cryostat and stored in 4% PFA in 1X PBS overnight. They were then transferred to a 30% sucrose + 1X PBS solution for cryoprotection. Free floating sections were washed 3x with 1X PBS and treated with a H_2_O_2_ solution (PBS + 10% methanol + 1.05% H_2_O_2_) for at least 1 h. Sections were washed, blocked for 2 h in 10% normal donkey serum + 1X PBS + 1% Triton X-100 (blocking solution) and then were treated overnight at 4°C with primary antibody diluted in blocking solution containing the following 2 antibodies: mouse monoclonal NET primary antibody, 1:200, Invitrotgen/Thermo Fisher Scientific, (Waltham, MA); rabbit polyclonal NR1 primary antibody, 1:100, Alomone Labs, (Jerusalem, Israel). The following morning, slices were washed and incubated for 2h at RT covered with secondary antibody diluted in blocking solution containing the following 2 antibodies: donkey anti-mouse Alexa Fluor 488, 1:2000, Invitrogen/Thermo Fisher Scientific, (Waltham, MA); donkey anti-rabbit Alexa Fluor 594, 1:1000, Invitrogen/Thermo Fisher Scientific, (Waltham, MA). Sections were next washed, mounted on gelatin covered slides and coverslipped using PermaFluor (Thermo Fisher Scientific, Waltham, MA) to preserve fluorescence for long-term storage at 4°C. Images were taken on a custom built 2-photon microscope at 20X magnification.

### Drugs

Where indicated, the following drugs were used: (-)-norepinephrine (NE; 20 μM), dopamine (DA; 20 μM), prazosin (α1-adrenergic antagonist; 2 μM), propranolol (β-adrenergic inhibitor; 5 μM), SCH 23390 (D1-like receptor antagonist; 10 µM), nisoxetine (norepinephrine transporter blocker; 5μM), RS 79948 (α2-adrenergic antagonist; 5 μM). All drugs were purchased from Tocris Bioscience (Minneapolis, MN).

### Statistical analysis

All electrophysiological data points are represented as the mean ± SEM. Field recordings were analyzed using two-way repeated measures ANOVAs with time as an independent variable with an assumption for a normal distribution at each averaged data point. Most ANOVAs were run over the last 30 minutes of recording after LTP stimulation. However, in Figure 4 an additional ANOVA was run over the 15 minutes that norepinephrine was present in panel A, and ANOVAs were run over the last 15 minutes of recording after LTP stimulation since recording only lasted 45 minutes after wLTP stimulation. Also, in Figure 2C another ANOVA was run over the first 30 minutes after LTP. All analyses were performed using GraphPad Prism 7 software (San Diego, CA).

## Data Availability

The data that support the findings of this study are available from the corresponding author upon reasonable request.

## Acknowledgements

We would like to thank Dr. Marc G. Caron for his generous gift of norepinephrine transporter knockout mice. We would also like to thank To Thai for his technical assistance.

## Author contributions

A.S. designed the experiments, performed the experiments, analyzed data, and wrote the paper.

R.W.G. designed the experiments and edited the paper.

## Additional information

The authors declare no competing interests.

---

**Supplementary Figure 1.**
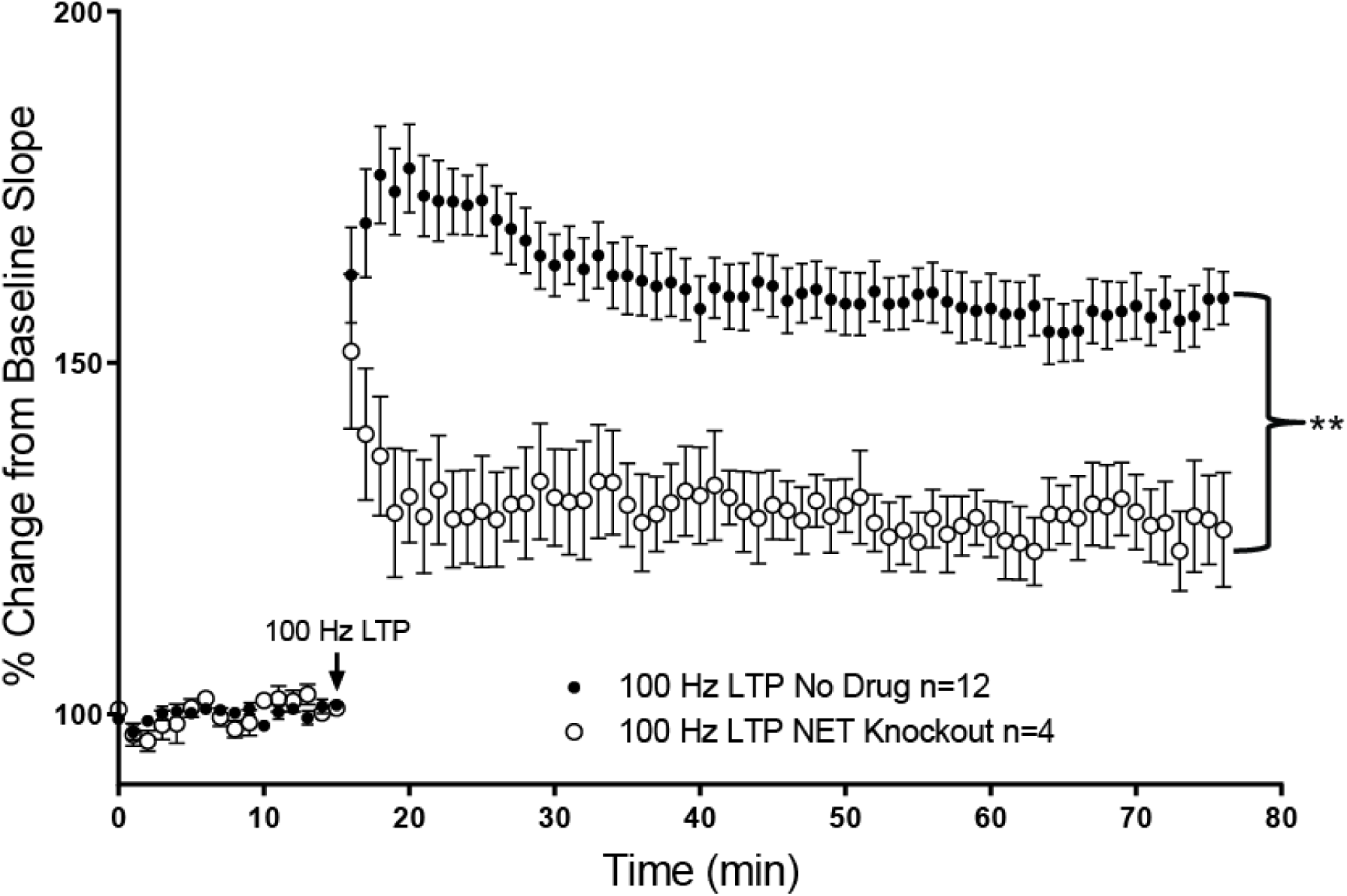
Knocking out the norepinephrine transporter (NET) reduces the magnitude of LTP in dorsal hippocampus. A different LTP protocol was administered (100 Hz for 1 second) without the addition of any adrenergic receptor antagonists in slices from Cre(-) control mice (black circles, n=12) and NET knockout mice (open circles, n=4). All data points are represented as mean +/- SEM. Tests for significance were done using a two-way repeated measures ANOVA over the last 30 minutes of 100 Hz LTP. Double asterisk represents p<0.01, F(1, 14) = 15.59, p=0.0015.

**Supplementary Figure 2.**
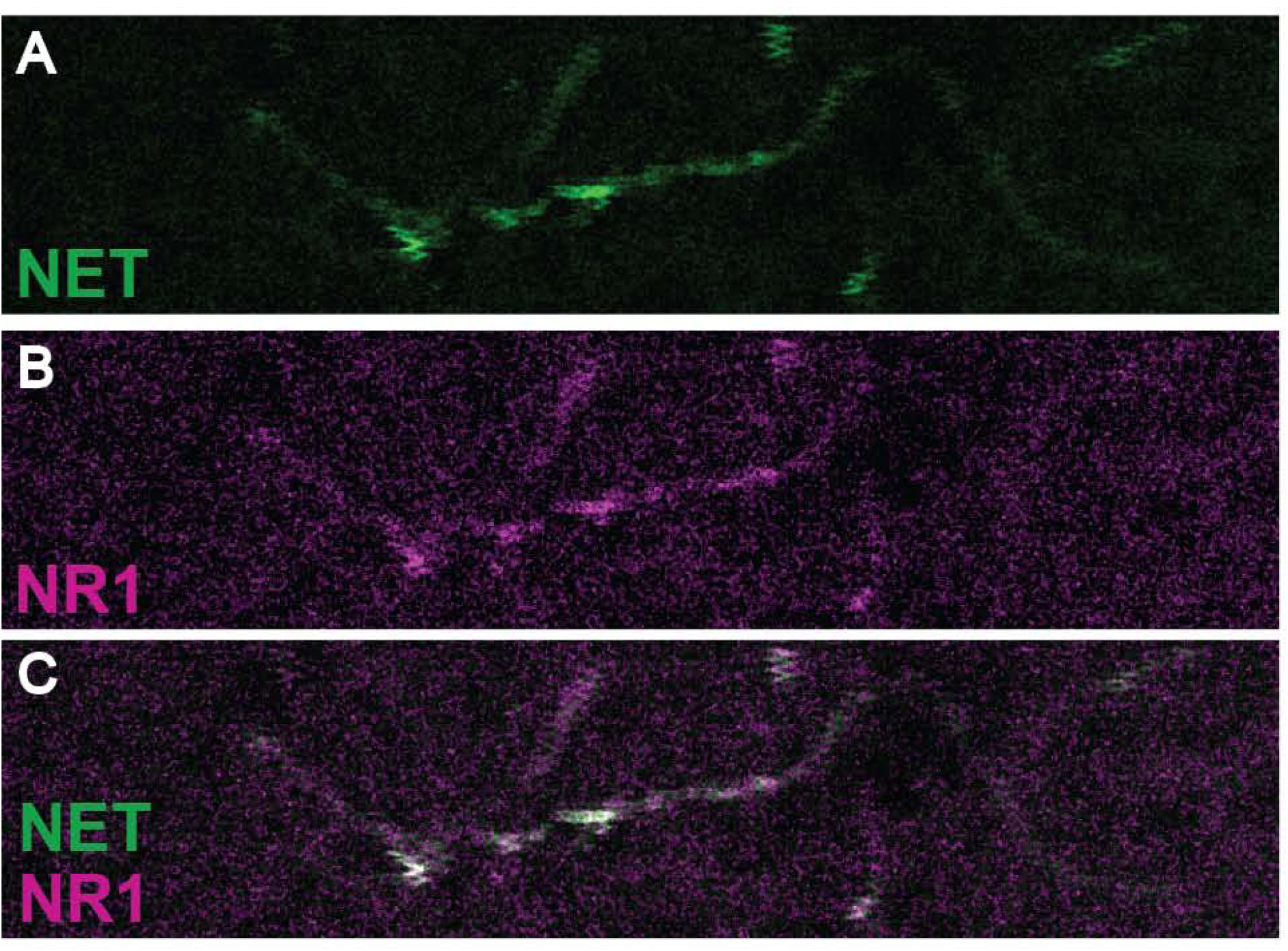
Colocalization of the norepinephrine transporter and presynaptic NMDA receptors in dorsal CA1. **A**,**B**,**C**, Representative immunnostaining image of LC fibers in the CA1 region of the dorsal hippocampus showing co-localization (**C**) of the norepinephrine transporter (NET, green, **A**) and presynaptic NMDA receptors (NR1, magenta, **B**) on LC terminals.

## References

1. Kebschull, J. M. et al. High-Throughput Mapping of Single-Neuron Projections by Sequencing of Barcoded RNA. Neuron 91, 975–987, doi:10.1016/j.neuron.2016.07.036 (2016).

2. Schwarz, L. A. et al. Viral-genetic tracing of the input-output organization of a central noradrenaline circuit. Nature 524, 88–92, doi:10.1038/nature14600 (2015).

3. Berridge, C. W. & Waterhouse, B. D. The locus coeruleus–noradrenergic system: modulation of behavioral state and state-dependent cognitive processes. Brain Research Reviews 42, 33–84, doi:10.1016/s0165-0173(03)00143-7 (2003).

4. Beas, B. S. et al. The locus coeruleus drives disinhibition in the midline thalamus via a dopaminergic mechanism. Nature neuroscience 21, 963–973, doi:10.1038/s41593-018-0167-4 (2018).

5. Kempadoo, K. A., Mosharov, E. V., Choi, S. J., Sulzer, D. & Kandel, E. R. Dopamine release from the locus coeruleus to the dorsal hippocampus promotes spatial learning and memory. Proceedings of the National Academy of Sciences of the United States of America 113, 14835–14840, doi:10.1073/pnas.1616515114 (2016).

6. Devoto, P., Flore, G., Saba, P., Fa, M. & Gessa, G. L. Stimulation of the locus coeruleus elicits noradrenaline and dopamine release in the medial prefrontal and parietal cortex. Journal of neurochemistry 92, 368–374, doi:10.1111/j.1471-4159.2004.02866.x (2005).

7. Zerbi, V. et al. Rapid Reconfiguration of the Functional Connectome after Chemogenetic Locus Coeruleus Activation. Neuron 103, 702–718 e705, doi:10.1016/j.neuron.2019.05.034 (2019).

8. Takeuchi, T. et al. Locus coeruleus and dopaminergic consolidation of everyday memory. Nature 537, 357–362, doi:10.1038/nature19325 (2016).

9. Wagatsuma, A. et al. Locus coeruleus input to hippocampal CA3 drives single-trial learning of a novel context. Proceedings of the National Academy of Sciences of the United States of America 115, E310–E316, doi:10.1073/pnas.1714082115 (2018).

10. Lemon, N. & Manahan-Vaughan, D. Dopamine D1/D5 receptors contribute to de novo hippocampal LTD mediated by novel spatial exploration or locus coeruleus activity. Cerebral cortex 22, 2131–2138, doi:10.1093/cercor/bhr297 (2012).

11. Lisman, J., Grace, A. A. & Duzel, E. A neoHebbian framework for episodic memory; role of dopamine-dependent late LTP. Trends in neurosciences 34, 536–547, doi:10.1016/j.tins.2011.07.006 (2011).

12. Nomura, S. et al. Noradrenalin and dopamine receptors both control cAMP-PKA signaling throughout the cerebral cortex. Frontiers in cellular neuroscience 8, 247, doi:10.3389/fncel.2014.00247 (2014).

13. Moron, J. A., Brockington, A., Wise, R. A., Rocha, B. A. & Hope, B. T. Dopamine uptake through the norepinephrine transporter in brain regions with low levels of the dopamine transporter: evidence from knock-out mouse lines. The Journal of neuroscience : the official journal of the Society for Neuroscience 22, 389–395 (2002).

14. Borgkvist, A., Malmlof, T., Feltmann, K., Lindskog, M. & Schilstrom, B. Dopamine in the hippocampus is cleared by the norepinephrine transporter. The international journal of neuropsychopharmacology / official scientific journal of the Collegium Internationale Neuropsychopharmacologicum 15, 531–540, doi:10.1017/S1461145711000812 (2012).

15. Robertson, S. D., Matthies, H. J. & Galli, A. A closer look at amphetamine-induced reverse transport and trafficking of the dopamine and norepinephrine transporters. Molecular neurobiology 39, 73–80, doi:10.1007/s12035-009-8053-4 (2009).

16. Smith, C. C. & Greene, R. W. CNS dopamine transmission mediated by noradrenergic innervation. The Journal of neuroscience : the official journal of the Society for Neuroscience 32, 6072–6080, doi:10.1523/JNEUROSCI.6486-11.2012 (2012).

17. Leviel, V. Dopamine release mediated by the dopamine transporter, facts and consequences. Journal of neurochemistry 118, 475–489, doi:10.1111/j.1471-4159.2011.07335.x (2011).

18. Khoshbouei, H., Wang, H., Lechleiter, J. D., Javitch, J. A. & Galli, A. Amphetamine-induced dopamine efflux. A voltage-sensitive and intracellular Na+-dependent mechanism. The Journal of biological chemistry 278, 12070–12077, doi:10.1074/jbc.M212815200 (2003).

19. Gnegy, M. E. et al. Intracellular Ca2+ regulates amphetamine-induced dopamine efflux and currents mediated by the human dopamine transporter. Molecular pharmacology 66, 137–143, doi:10.1124/mol.66.1.137 (2004).

20. Olivier, V., Guibert, B. & Leviel, V. Direct in vivo comparison of two mechanisms releasing dopamine in the rat striatum. Brain research 695, 1–9, doi:10.1016/0006-8993(95)00706-v (1995).

21. Falkenburger, B. H., Barstow, K. L. & Mintz, I. M. Dendrodendritic inhibition through reversal of dopamine transport. Science 293, 2465–2470, doi:10.1126/science.1060645 (2001).

22. Bermingham, D. P. & Blakely, R. D. Kinase-dependent Regulation of Monoamine Neurotransmitter Transporters. Pharmacol Rev 68, 888–953, doi:10.1124/pr.115.012260 (2016).

23. Andersen, J., Ringsted, K. B., Bang-Andersen, B., Stromgaard, K. & Kristensen, A. S. Binding site residues control inhibitor selectivity in the human norepinephrine transporter but not in the human dopamine transporter. Scientific reports 5, 15650, doi:10.1038/srep15650 (2015).

24. Kwon, O. B. et al. Neuregulin-1 regulates LTP at CA1 hippocampal synapses through activation of dopamine D4 receptors. Proceedings of the National Academy of Sciences of the United States of America 105, 15587–15592, doi:10.1073/pnas.0805722105 (2008).

25. Mather, M., Clewett, D., Sakaki, M. & Harley, C. W. Norepinephrine ignites local hotspots of neuronal excitation: How arousal amplifies selectivity in perception and memory. The Behavioral and brain sciences 39, e200, doi:10.1017/S0140525X15000667 (2016).

26. Hires, S. A., Zhu, Y. & Tsien, R. Y. Optical measurement of synaptic glutamate spillover and reuptake by linker optimized glutamate-sensitive fluorescent reporters. Proc Natl Acad Sci U S A 105, 4411–4416, doi:10.1073/pnas.0712008105 (2008).

27. Chaki, S., Okuyama, S., Ogawa, S. & Tomisawa, K. Regulation of NMDA-induced [3H]dopamine release from rat hippocampal slices through sigma-1 binding sites. Neurochemistry international 33, 29–34 (1998).

28. Malva, J. O., Carvalho, A. P. & Carvalho, C. M. Modulation of dopamine and noradrenaline release and of intracellular Ca2+ concentration by presynaptic glutamate receptors in hippocampus. British journal of pharmacology 113, 1439–1447, doi:10.1111/j.1476-5381.1994.tb17158.x (1994).

29. Lawrence, J. J. & Cobb, S. in Hippocampal Microcircuits 227–325 (Springer, 2018).

30. Jenson, D. et al. Dopamine and norepinephrine receptors participate in methylphenidate enhancement of in vivo hippocampal synaptic plasticity. Neuropharmacology 90, 23–32, doi:10.1016/j.neuropharm.2014.10.029 (2015).

31. Ramos, B. P. & Arnsten, A. F. Adrenergic pharmacology and cognition: focus on the prefrontal cortex. Pharmacology & therapeutics 113, 523–536, doi:10.1016/j.pharmthera.2006.11.006 (2007).

32. Nguyen, P. V. & Kandel, E. R. Brief theta-burst stimulation induces a transcription-dependent late phase of LTP requiring cAMP in area CA1 of the mouse hippocampus. Learning & memory 4, 230–243 (1997).

33. Larson, J. & Munkacsy, E. Theta-burst LTP. Brain research 1621, 38–50, doi:10.1016/j.brainres.2014.10.034 (2015).

34. Abercrombie, E. D., Keller, R. W., Jr. & Zigmond, M. J. Characterization of hippocampal norepinephrine release as measured by microdialysis perfusion: pharmacological and behavioral studies. Neuroscience 27, 897–904 (1988).

35. Hagena, H., Hansen, N. & Manahan-Vaughan, D. beta-Adrenergic Control of Hippocampal Function: Subserving the Choreography of Synaptic Information Storage and Memory. Cerebral cortex 26, 1349–1364, doi:10.1093/cercor/bhv330 (2016).

36. Raiteri, M., Garrone, B. & Pittaluga, A. N-methyl-D-aspartic acid (NMDA) and non-NMDA receptors regulating hippocampal norepinephrine release. II. Evidence for functional cooperation and for coexistence on the same axon terminal. The Journal of pharmacology and experimental therapeutics 260, 238–242 (1992).

37. Matsushita, N. et al. Dynamics of tyrosine hydroxylase promoter activity during midbrain dopaminergic neuron development. Journal of neurochemistry 82, 295–304, doi:10.1046/j.1471-4159.2002.00972.x (2002).

38. Palacios-Filardo, J. & Mellor, J. R. Neuromodulation of hippocampal long-term synaptic plasticity. Current opinion in neurobiology 54, 37–43, doi:10.1016/j.conb.2018.08.009 (2019).

39. Harley, C. W. Norepinephrine and dopamine as learning signals. Neural plasticity 11, 191–204 (2004).

40. Huang, Y. Y. & Kandel, E. R. D1/D5 receptor agonists induce a protein synthesis-dependent late potentiation in the CA1 region of the hippocampus. Proceedings of the National Academy of Sciences of the United States of America 92, 2446–2450 (1995).

41. O’Carroll, C. M., Martin, S. J., Sandin, J., Frenguelli, B. & Morris, R. G. Dopaminergic modulation of the persistence of one-trial hippocampus-dependent memory. Learning & memory 13, 760–769, doi:10.1101/lm.321006 (2006).

42. O’Dell, T. J., Connor, S. A., Gelinas, J. N. & Nguyen, P. V. Viagra for your synapses: Enhancement of hippocampal long-term potentiation by activation of beta-adrenergic receptors. Cellular signalling 22, 728–736, doi:10.1016/j.cellsig.2009.12.004 (2010).

43. Katsuki, H., Izumi, Y. & Zorumski, C. F. Noradrenergic regulation of synaptic plasticity in the hippocampal CA1 region. Journal of neurophysiology 77, 3013–3020, doi:10.1152/jn.1997.77.6.3013 (1997).

44. Bergles, D. E., Doze, V. A., Madison, D. V. & Smith, S. J. Excitatory actions of norepinephrine on multiple classes of hippocampal CA1 interneurons. The Journal of neuroscience : the official journal of the Society for Neuroscience 16, 572–585 (1996).

45. Rosen, Z. B., Cheung, S. & Siegelbaum, S. A. Midbrain dopamine neurons bidirectionally regulate CA3-CA1 synaptic drive. Nature neuroscience 18, 1763–1771, doi:10.1038/nn.4152 (2015).

46. Otmakhova, N. A. & Lisman, J. E. Dopamine selectively inhibits the direct cortical pathway to the CA1 hippocampal region. The Journal of neuroscience : the official journal of the Society for Neuroscience 19, 1437–1445 (1999).

47. Ito, H. T. & Schuman, E. M. Frequency-dependent gating of synaptic transmission and plasticity by dopamine. Frontiers in neural circuits 1, 1, doi:10.3389/neuro.04.001.2007 (2007).

48. Chiti, Z. & Teschemacher, A. G. Exocytosis of norepinephrine at axon varicosities and neuronal cell bodies in the rat brain. FASEB journal : official publication of the Federation of American Societies for Experimental Biology 21, 2540–2550, doi:10.1096/fj.06-7342com (2007).

49. Chun, M. M. & Turk-Browne, N. B. Interactions between attention and memory. Current opinion in neurobiology 17, 177–184, doi:10.1016/j.conb.2007.03.005 (2007).

50. Sara, S. J. The locus coeruleus and noradrenergic modulation of cognition. Nature reviews. Neuroscience 10, 211–223, doi:10.1038/nrn2573 (2009).

51. Devilbiss, D. M. Consequences of tuning network function by tonic and phasic locus coeruleus output and stress: Regulating detection and discrimination of peripheral stimuli. Brain Res 1709, 16–27, doi:10.1016/j.brainres.2018.06.015 (2019).

52. Berridge, C. W. & Abercrombie, E. D. Relationship between locus coeruleus discharge rates and rates of norepinephrine release within neocortex as assessed by in vivo microdialysis. Neuroscience 93, 1263–1270 (1999).

53. Aston-Jones, G. & Cohen, J. D. An integrative theory of locus coeruleus-norepinephrine function: adaptive gain and optimal performance. Annual review of neuroscience 28, 403–450, doi:10.1146/annurev.neuro.28.061604.135709 (2005).

54. Fog, J. U. et al. Calmodulin kinase II interacts with the dopamine transporter C terminus to regulate amphetamine-induced reverse transport. Neuron 51, 417–429, doi:10.1016/j.neuron.2006.06.028 (2006).

55. Foster, J. D. & Vaughan, R. A. Phosphorylation mechanisms in dopamine transporter regulation. J Chem Neuroanat 83-84, 10–18, doi:10.1016/j.jchemneu.2016.10.004 (2017).

56. Aggarwal, S. & Mortensen, O. V. Overview of Monoamine Transporters. Current protocols in pharmacology 79, 12 16 11–12 16 17, doi:10.1002/cpph.32 (2017).

57. Redondo, R. L. & Morris, R. G. Making memories last: the synaptic tagging and capture hypothesis. Nat Rev Neurosci 12, 17–30, doi:10.1038/nrn2963 (2011).

58. Mlinar, B., Stocca, G. & Corradetti, R. Endogenous serotonin facilitates hippocampal long-term potentiation at CA3/CA1 synapses. Journal of neural transmission 122, 177–185, doi:10.1007/s00702-014-1246-7 (2015).

59. Thompson, A. M., Swant, J. & Wagner, J. J. Cocaine-induced modulation of long-term potentiation in the CA1 region of rat hippocampus. Neuropharmacology 49, 185–194, doi:10.1016/j.neuropharm.2005.03.005 (2005).

60. Patriarchi, T. et al. Ultrafast neuronal imaging of dopamine dynamics with designed genetically encoded sensors. Science 360, doi:10.1126/science.aat4422 (2018).

61. Sun, F. et al. A Genetically Encoded Fluorescent Sensor Enables Rapid and Specific Detection of Dopamine in Flies, Fish, and Mice. Cell 174, 481–496 e419, doi:10.1016/j.cell.2018.06.042 (2018).

62. Feng, J. et al. A Genetically Encoded Fluorescent Sensor for Rapid and Specific In Vivo Detection of Norepinephrine. Neuron 102, 745–761 e748, doi:10.1016/j.neuron.2019.02.037 (2019).

